# Context triggers the retrieval of long-term memories into working memory to guide attention

**DOI:** 10.1101/401216

**Authors:** Sirawaj Itthipuripat, Geoffrey F Woodman

**Author notes:** Correspondence Sirawaj Itthipuripat or Geoffrey F Woodman, or, Vanderbilt University, Department of Psychology, 111 21st Ave South, Nashville, TN 37240.

## Abstract

How do we know what we are looking for in familiar scenes and surroundings? Here we tested a novel hypothesis derived from theories of human memory that working memory (WM) buffers mnemonic contents retrieved from long-term memory (LTM) to control attention. To test this hypothesis, we measured the electrical fields recorded noninvasively from human subjects’ as they searched for specific sets of objects in learned contexts. We found that the subjects’ WM-indexing brain activity tracked the number of real-world objects people learned to search for in each context. Moreover, the level of this WM activity predicted the inter-subject variability in behavioral performance. Together, our results demonstrate that familiar contexts can trigger the transfer of information from LTM to WM to provide top-down attentional control.

## Introduction

When an experienced driver gets behind the wheel of an automobile, how is it that they can immediate shift attention to task-relevant information (e.g., red break lights, pedestrians entering the road, trucks looming in the rearview mirror)? Theories of human memory have suggested that context can be used to pull mnemonic information out of LTM and into working memory (WM) (Atkinson and Shiffrin, 1968; Broadbent, 1975; Cowan, 2001; Fukuda and Woodman, 2017; James, 1890). However, theories of attention have never considered how this kind of explicit memory retrieval in well-learned contexts might allow us to control which objects we direct attention to (Bundesen et al., 2005; Desimone and Duncan, 1995; Wolfe and Horowitz, 2004; Woodman et al., 2013). Past studies have shown that repeated exposure to specific objects in specific contexts can facilitate attentional selection and eye movements to task-relevant targets and this engages brain areas in the medial temporal lobe including the hippocampus (Chun and Jiang, 1998; Chun and Phelps, 1999; Giesbrecht et al., 2013; Goldfarb et al., 2016; Greene et al., 2007; Hannula and Ranganath, 2009; Preston and Gabrieli, 2008; Summerfield et al., 2006). But how this learning influences how attention is controlled is still not well understood, with a number of accounts proposing that learning can influence attention implicitly, without any storage of information in WM (Chun and Jiang, 1998; Chun and Phelps, 1999). According to this account, the LTM representations can directly influence how attention is deployed to objects, without them needing to be buffered in WM. Here we aimed to test the novel hypothesis derived from theories of memory in which representations retrieved from LTM are actively maintained in WM to provide top-down control over attentional selection.

If it is true that learned context can trigger the retrieval of LTMs into WM to guide attention, we should observe a neural index of WM maintenance as people re-encounter a familiar context. To test this hypothesis, we recorded the electroencephalogram (EEG) from human subjects who performed a visual search task where they searched for specific targets in specific contexts that varied in their background color (Figure 1a). To infer whether LTM representations were being buffered in WM, we measured the subjects’ posterior delay activity (pDA). We predicted that the amplitude of this posterior WM-related activity would increase as function of the number of targets previously paired with a given context, up until subjects’ WM was filled with possible targets (Brady et al., 2016; Cowan, 2001; Luck et al., 1997; Todd and Marois, 2004; Vogel and Machizawa, 2004; Vogel et al., 2005; Xu and Chun, 2006). In addition, if our index of WM storage of the searched-for targets was the driver of subjects’ performance, then we should see that the amplitude of this activity is related to subjects’ search performance.

**Figure 1.**
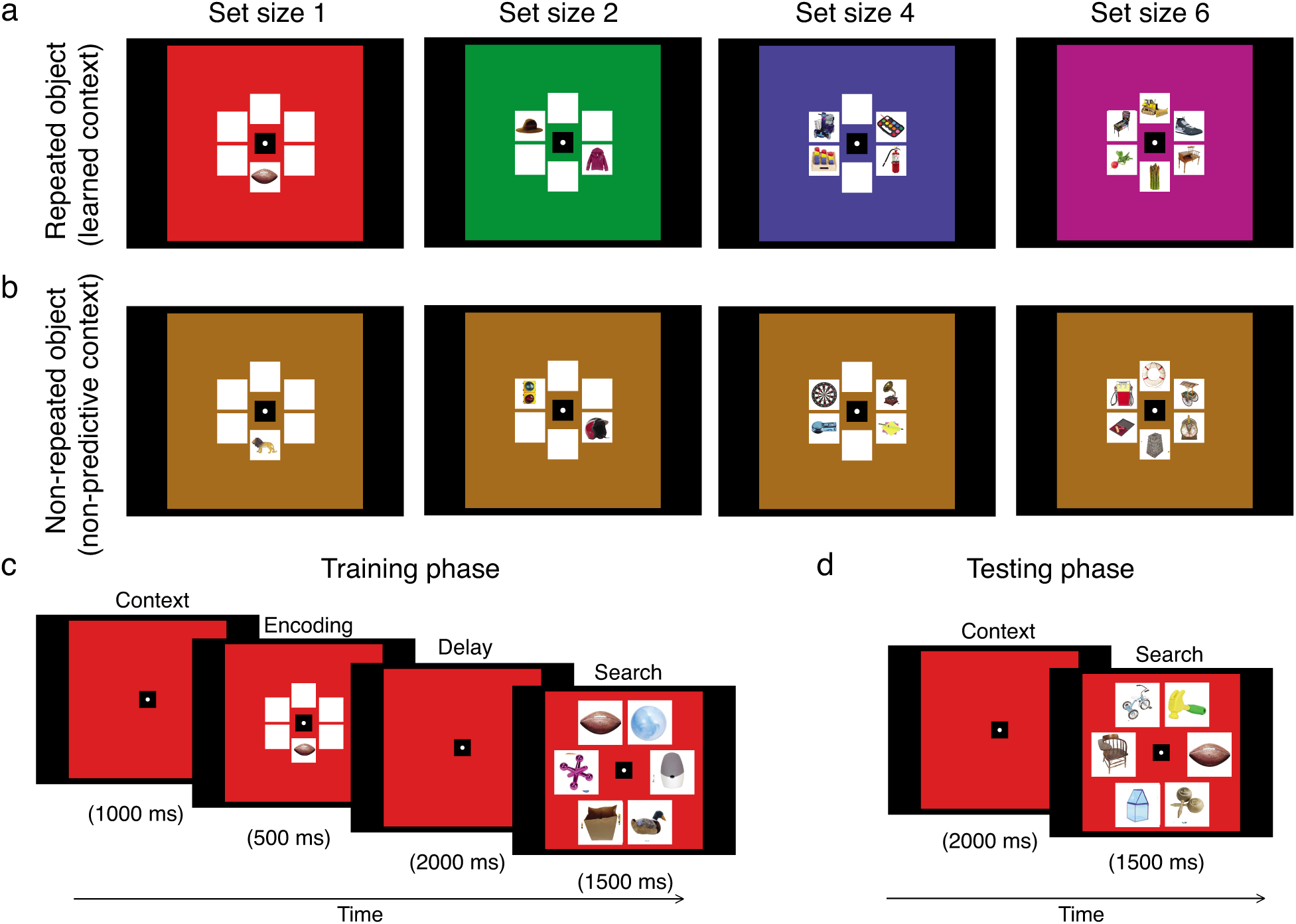
Task and behavioral results. (a) Human subjects (*N* = 36) learned to find 4 different sets of unique real-world objects within 4 different colored background contexts (b) They also encountered another colored background followed by objects that never repeated. (c-d) Examples for trial structures during the training and test phases, respectively.

## Results

Thirty-six female and male human subjects performed a visual search task where they had to search for specific targets in different colored background contexts (Figure 1a). During training, these subjects repeatedly searched for a specific set of objects in each of 4 different contexts for 120 trials. As a comparison condition, we interleaved trials with a fifth background context color in which the searched for targets were unique on each trial (i.e., non-predictive context; Figure 1b). Across other 120 trials, this non-predictive context was associated with 390 unique objects (i.e., each of 30 from 120 trials contained 1, 2, 4, and 6 non-repeated objects). This condition was the most difficult, but LTM representations of the set of possible targets could not be retrieved to prepare for the search array. Thus, this provides us with a control condition that rules out explanations based on simple task difficulty that increases with the increasing number of memory items. Each training trial started with the presentation of the background color alone, followed by an array of 1, 2, 4, or 6 unique real-world objects (Brady et al., 2008) indicating the target or targets that subjects were to search for following a blank delay of 2000ms (Figure 1c). Subjects were instructed to learn the associations between different color contexts and those targets. Then, in the final test phase, they had to search for the possible targets in each context without seeing those targets (Figure 1d). Thus, if context triggers the retrieval of LTM information, which is then buffered by WM, the onset of the context should elicit WM-related activity as subjects fill WM with the possible targets given that context.

As shown in Figure 2 left, in the training phase, subjects learned to detect the targets in each color context. While there was a general decrease in target sensitivity (d’) with increasing set size, training significantly improved d’ and the training effect was larger for the larger set size (compare filled circles to filled squares in Figure 2 top right, trial repetition effect: (F(1, 35) = 133.33, p = 1.73 × 10^−13^; set size × trial repetition interaction: F(3, 105) = 8.44, p = 4.89 × 10^−5^). Reaction times from correct trials (correct RTs) also increased as a function of set size (F(3, 105) = 269.72, p ≤ 1.00 × 10^−15^) but did not change significantly across the training phase (compare filled circles to squares in Figure 2 bottom right, F(1, 35) = 2.26, p = 0.142).

**Figure 2.**
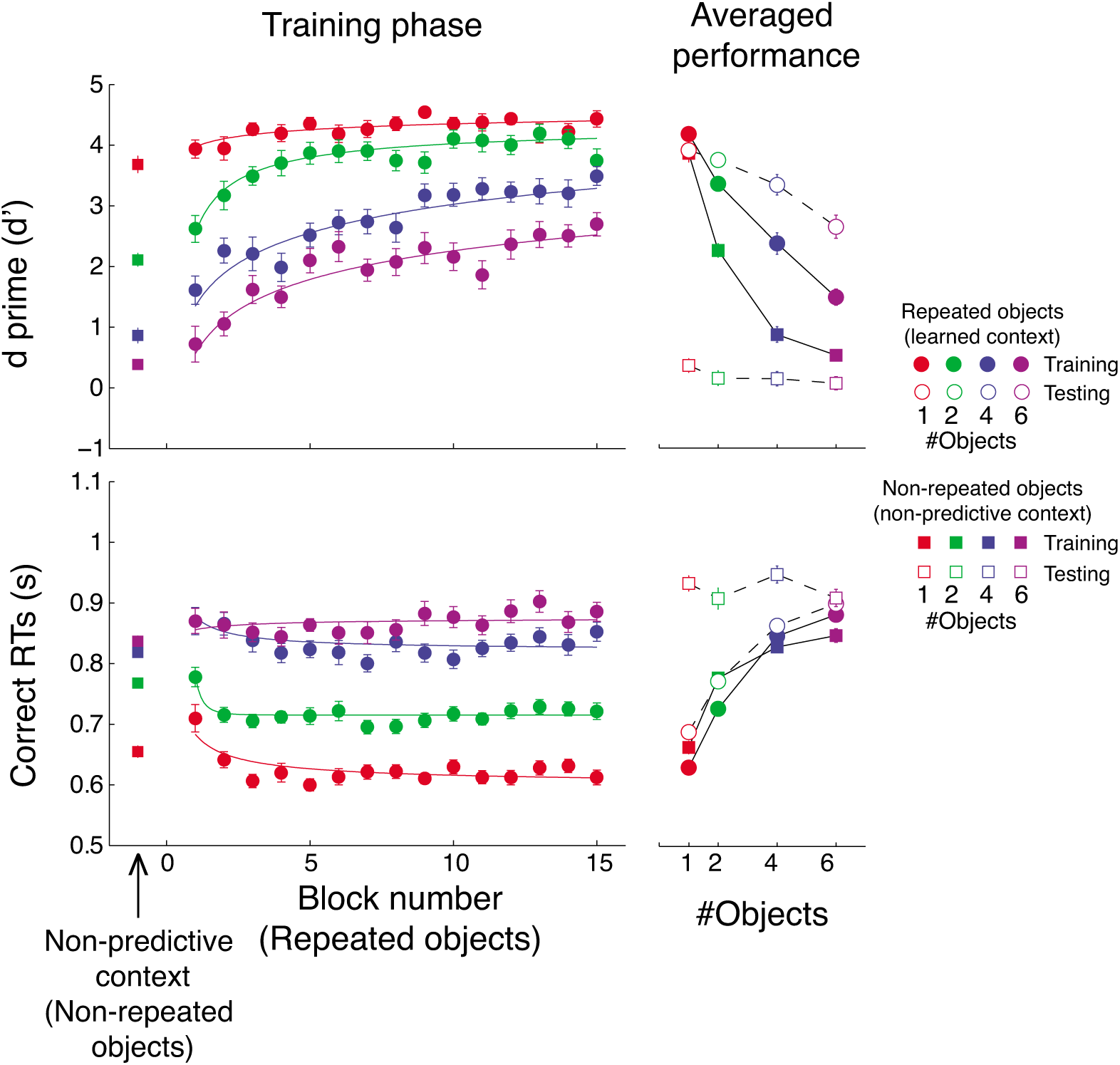
Behavioral results. (Top) search d-prime (d’). (Bottom) RTs on correct trials. (Left) Learning performance across blocks of trials during the training phase. The data were fit with a power function. (Right) Averaged performance for repeated objects (learned context) and non-repeated objects (non-predictive-context) plotted as function of set size during both training and test phases. Error bars represent ±1 within-subject SEM.

During the test phase, d’ decreased as set size increased (F(3, 105) = 55.02, p ≤ 1.00 × 10^−15^), but the set size effect was relatively small compared to that of the training phase as training compressed performance at ceiling levels (compared empty to filled circles in Figure 2 top right, phase effect: F(1, 35) = 47.40, p = 5.39 × 10^−8^; set size × phase interaction: F(3, 105) = 22.28, p = 3.07 × 10^−11^). While d’ for all learned contexts was near ceiling after training, d’ for the non-predictive context was near zero (compared empty circle to empty squares in Figure 2 top right, repetition effect: F(1, 35) = 480.13, p ≤ 1.00 × 10^−15^). This near-chance performance for the non-predictive context was expected because of the minimally training with any one of the many different possible targets (i.e., subjects saw each of 390 targets only once during training). Overall, correct RTs for repeated contexts during the test phase also increased as a function of set size (F(3, 105) = 92.03, p ≤ 1.00 × 10^−15^), and they were slightly longer during the test phase than the training phase (compared empty circle to empty squares in Figure 2 bottom right, phase effect: F(1, 35) = 19.18, p = 1.03 × 10^−4^). A further analysis showed that this was true only for correct rejection RTs (F(1, 35) = 38.86, p = 3.80 × 10^−7^) but not for hit RTs (F(1, 35) = 0.08, p = 0.782). This suggests that the subjects may have been more cautious to reject target-absent trials when they had to pull information from LTM. Similar to the d’ result, during the test phase, correct RTs were much shorter for all learned target contexts compared to those in trials with the non-predictive context (compared empty circle to empty squares in Figure 2 top right, repetition effect: F(1, 35) = 56.73, p ≤ 1.00 × 10^−15^).

If humans use context to retrieve mnemonic contents from LTM and buffer the retrieved information in WM before performing the search task, we should observe a neural index of visual WM maintenance as people re-encounter a familiar context. Specifically, in the test phase, we should see the background color elicits a pDA component with an amplitude corresponding to the number of possible targets (Fukuda et al., 2015), or the capacity of WM if the number of possible targets exceeds the capacity limit (i.e., up until visual WM capacity of 3-4 memory items) (Brady et al., 2016; Cowan, 2001; Luck et al., 1997; Todd and Marois, 2004; Vogel and Machizawa, 2004; Vogel et al., 2005; Xu and Chun, 2006).

We found a significant main effect of set size on pDA amplitude from 400-900ms after the cue onset (Figures 3a.i-iii, F(3, 105) = 7.41, p =1.50 × 10^−4^,). The pDA amplitude increased (i.e., became more negative) as the number of to-be-retrieved targets increased, and saturated at set size 4 (see post-hoc t-test results in Supplementary Table 1). Importantly, when the number of to-be-retrieved targets was 390 (i.e., the non-predictive context), the pDA amplitude returned to a baseline activity level that did not significantly differ from set size 1 (planned paired t-tests showed significant differences between the non-predictive context and the learned contexts paired with 2, 4, and 6 targets: t’s(35) = −5.83 to −2.37, p’s = 1.29 × 10^−6^ to 0.0236). This confirms that the pDA amplitude does not simply index general changes in task difficulty, but actual LTM contents because LTM representations in the non-predictive context were not accessible to aid search (i.e., search d’ = ∼0, see Figure 2). An auxiliary analysis showed that the set size effect on the pDA amplitude was significant only in subjects who did well learning, but not in subjects who poorly learned the targets (Supplementary Figure 1). Furthermore, the pDA closely tracked inter-subject variability in search performance during the test phase. Specifically, behavioral d’ during the test phase was significantly correlated with the amplitude of the pDA for set sizes 4 and 6, normalized by the magnitude of the pDA measured from the non-predictive context (Figure 3a.iii, Rho’s = −0.47 and −0.35, p’s= −0.0034 and 0.038 for set sizes 4 and 6, respectively). This correlation suggests that LTM retrieval is limited by each subject’s capacity to buffer items in WM, emphasizing the tight relationship between the two memory systems and their essential roles in retrieving mnemonic information to guide attention.

**Figure 3.**
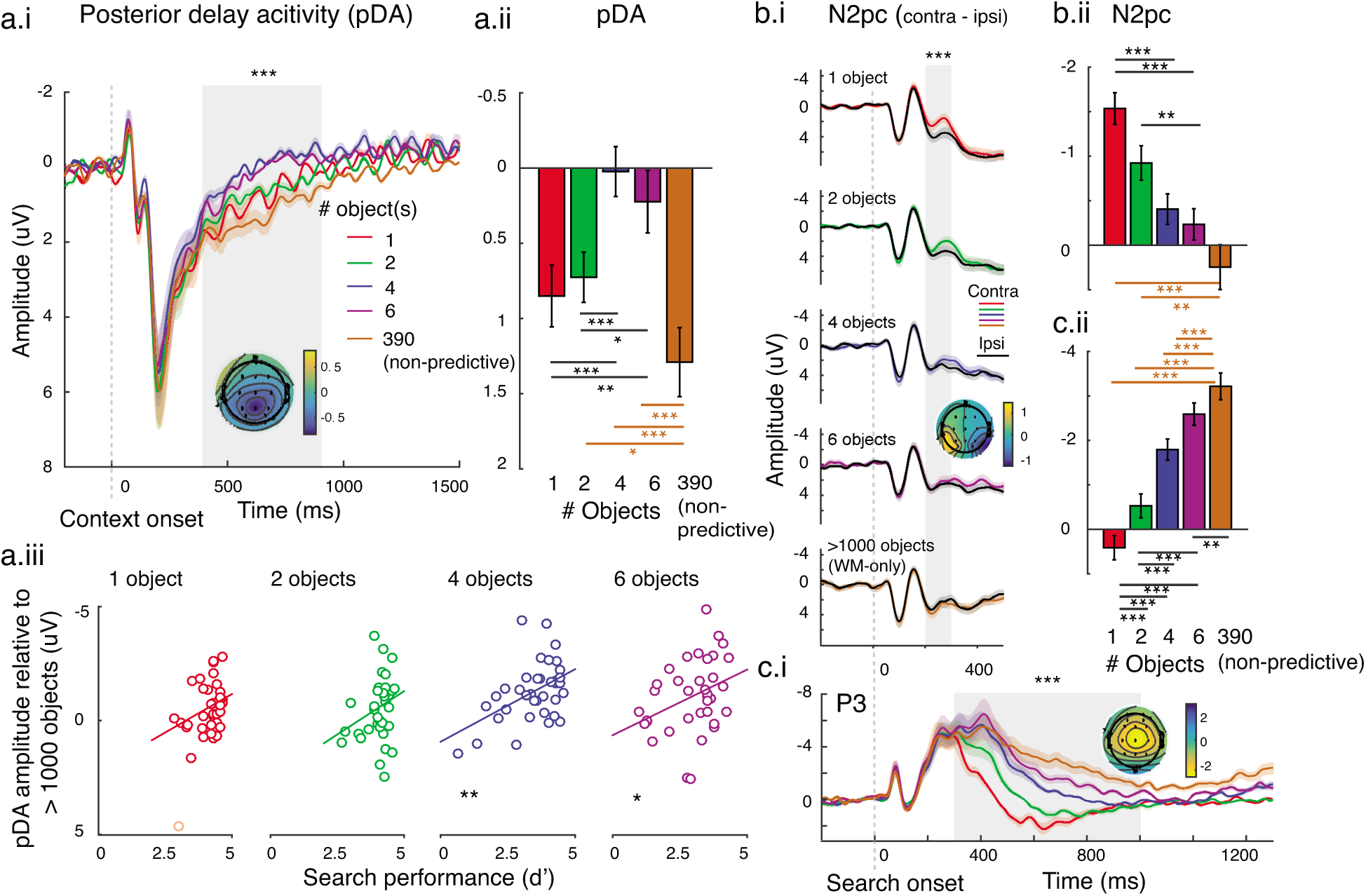
Neurophysiological results showing that learned context triggers WM activity indexing the number of the possible targets to guide attention and decision-making. (a.i-ii) The posterior delay negativity (pDA) tracked the number of targets possible in that context, up until WM capacity was reached. (a.iii) pDA amplitude relative to the non-predictive-context control condition correlated with search d-prime. (b.i-ii) The N2pc component on hit trials showed earlier and less variable shifts of spatial attention to the targets with fewer retrieved items. (c.i-ii) The P3 component was also larger in amplitude when fewer targets were possible. Both N2pc and P3 results were consistent with set-size effects on RTs. *** in a.i, b.i, and c.i show significant set size effects for the EEG data averaged across the shaded areas via repeated-measured one-way ANOVAs with p’s <0.001. Black *, **, and *** in Figures a.ii, b.ii, and c.ii show significant differences from post-hoc t-tests with p < 0.05, p < 0.01, and p < 0.001, respectively (Holm-Bonferroni corrected). Brown *, **, and *** in Figures a.ii, b.ii, and c.ii show significant differences from planned t-tests between non-predictive-context trials and learn context conditions with p < 0.05, p < 0.01, and p < 0.001, respectively. * and ** in a.iii show significant correlations with p <0.05 and p<0.001, respectively. A faint red dot in Figure a.iii (first panel) indicates an outlier excluded from the correlation analysis. All shaded areas around ERP traces and all error bars represent ±1 within-subject SEM. Note that we plotted negative up on all y-axes by electrophysiological convention.

We also examined the set size effect on the posterior-occipital alpha suppression (i.e., changes in power of EEG oscillations at 8-13Hz). Previous research suggests that alpha can track the storage of information in WM, even when it is retrieved from LTM (Foster et al., 2017b, 2016, Fukuda et al., 2015, 2016). Unlike the pDA, the alpha suppression during the delay period did not change as a function of memory set size (see all alpha results and statistics in Supplementary Figure 2). After the search onset, the magnitude of alpha suppression increased as a function of set size. However, the magnitude of alpha suppression in the non-predictive context was similar to target set sizes 4 and 6. This fits with the notion that modulations of alpha oscillations reflect the storage of purely spatial information, as well as general task demands, but with alpha activity containing no feature-or object-based information (Bae and Luck, 2017; Bosman et al., 2012; Foster et al., 2017b, 2017a, 2016; Foxe and Snyder, 2011; Foxe et al., 1998; Fries, 2001; Fries et al., 2008; Fukuda and Woodman, 2017; Klimesch et al., 2007; van Moorselaar et al., 2017; Nelli et al., 2017; Rihs et al., 2007; Rungratsameetaweemana et al., 2018; Sauseng et al., 2005; von Stein et al., 2000; de Vries et al., 2017).

Finally, we tested how the retrieved information could help guide attention and decision-making during the actual period in which search was conducted. Following previous studies, we used the N2pc (200-300ms) and the P3 components (300-900ms) to index the deployment of spatial attention to the target location and decision-making processes, respectively (Cosman et al., 2016; Eimer, 1996; Hickey et al., 2010; Hillyard et al., 1972; Itthipuripat et al., 2014, 2015, 2017; Squires et al., 1975; Woodman and Luck, 1999; Woodman et al., 2009). The amplitudes of the N2pc and the P3 decreased as a function of memory set size (Figures 3b-c; F(3,135)’s = 9.15 and 43.48 with p’s = 1.97 × 10^−5^ and ≤ 1.00 × 10^−15^ for N2pc and P300, respectively; also see post-hoc t-test results in Supplementary Table 1). In addition, the mean amplitudes of the N2pc and P3 were the lowest in the non-predictive context. Specifically, for the N2pc, planned t-tests show significant differences between non-predictive-context trials and trials with memory set sizes 1 and 2 (t(35)’s = −5.08 and −3.39, p’s = 1.27 × 10^−5^and 0.0017, respectively) but no differences between non-predictive-context trials and trials with memory set sizes 4 and 6 (t(35)’s = −1.83 and −1.28, p’s = 0.0760 and 0.2106, respectively). For the P3, planned t-tests show significant differences between non-predictive-context trials and all other set sizes (t(35)’s = 2.31-10.25, p’s = 0.0269 – 4.43 × 10^−12^). These modulations of the N2pc and P3 components are consistent with the set size effects on search RT.

## Discussion

Here we show that WM-related activity (i.e., the pDA component) can track the retrieval of target representations from LTM into WM that occurs spontaneously when people view a context in which they learned to search for specific sets of targets. As we predicted by deriving a novel hypothesis from cognitive theories of human memory, the pDA amplitude tracked the number of retrieved LTM representations up until WM was filled (i.e., at ∼3-4 memory items) (Atkinson and Shiffrin, 1968; Broadbent, 1975; Cowan, 2001; Fukuda and Woodman, 2017; James, 1890). Moreover, the magnitude of the pDA modulation explained inter-subject variability in behavioral performance during the test phase.

Our results are highly similar to results from previous studies that measure the parietal activity in WM tasks, where subjects maintain new incoming sensory inputs (Brady et al., 2016; Cowan, 2001; Luck et al., 1997; Todd and Marois, 2004; Vogel and Machizawa, 2004; Vogel et al., 2005; Xu and Chun, 2006). In those studies, the parietal activity scaled up with the number of sensory inputs and saturated at set sizes 3-4, which is thought to be the maximum capacity of visual WM (Brady et al., 2016; Cowan, 2001; Fukuda et al., 2015; Luck et al., 1997; Todd and Marois, 2004; Vogel and Machizawa, 2004; Vogel et al., 2005; Xu and Chun, 2006). Moreover, the level of activity also correlated with WM capacity of individual subjects estimated with different behavioral markers (Fukuda et al., 2015; Vogel and Machizawa, 2004), and manipulated by electrical stimulation over the parietal lobe (Heinen et al., 2016; Li et al., 2017; Tseng et al., 2012). Together, the similarity between our pDA results and those reported by previous WM studies suggest the parietal activity plays a common role in actively maintaining new incoming sensory information and also LTM representations as they are retrieved back into WM. Moreover, our findings suggest that WM is a key mechanism by which top-down attentional control is maintained even after extensive learning. Together, these help extend theories of attention, which have not previously explained how top-down control settings in different environmental contexts are acquired and maintained across time (Bundesen et al., 2005; Desimone and Duncan, 1995; Wolfe and Horowitz, 2004; Woodman et al., 2013).

Our observations help knit together proposals from theories of attention with proposals from theories of human memory. Specifically, theories of top-down attentional control have proposed that WM representations have a special status in being able to control attentional selection (Olivers et al., 2011), and our findings demonstrate that representations are held in WM even in well-learned environments, a process that could take advantage of this special status. What we do not yet know is whether WM is used in all situations. For example, it is possible for people to search for hundreds of possible targets at the same time (Wolfe, 2012), and this may rely exclusively on LTM representations that do not need to be held in WM. Alternatively, it is possible that these situations with huge possible target sets involve maintaining prototype-like representations in WM for entire categories of objects (i.e., any bird), with this circumventing the capacity limit inherent in WM storage. Future experiments using the method that we introduce here will be able to address questions such as these regarding the ubiquity of the phenomenon we discovered here.

## Acknowledgements

We thank Paul Shin and Brittany L. Correia for helping with data collection. We also thank Adam R Aron, Jason Rajsic, Sisi Wang, Christopher Sundby, and Mathieu Servant for technical help and useful discussions.

## Author contributions

SI conceived and implemented the experiment, collected and analyzed the data, wrote the first draft of the manuscript, and edited the manuscript. GFW conceived and supervised the project, and co-wrote the manuscript.

## Materials and Methods

### Subjects

We recruited 40 neurologically healthy male and female adult humans, who had normal or corrected-to-normal color vision, from the Vanderbilt University (VU) and Nashville community. All subjects provided written informed consent as required by the local Institutional Review Board at VU (IRB#071176). They were compensated at a rate of $10 USD per hour of participation. Four subjects did not complete the experimental protocol, leaving 36 subjects in the final analysis (age range = 18-39 years old, mean age = 23.31±5.48 SD, 28 male, 4 left-handed). We collected a relatively large sample of data because we aimed to investigate individual differences in retrieval performance and neural indexes of WM (see below).

### Stimuli and Tasks

Stimuli were presented on a Macintosh desktop running MATLAB 7.10.0 (R2010A) (Mathworks Inc., Natick, MA) and the Psychophysics Toolbox (version 3.0.8(Brainard, 1997; Pelli, 1997)) Subjects were seated 80 cm from the CRT monitor with a black background of 1.23 cd/m^2^(refresh rate = 75 Hz, 800 × 600 resolution). The experiment was performed in a dark, sound-attenuated, and electromagnetically shielded room (ETS-Lindgren, Cedar Park, TX, USA). The real-world object images used in the experiment were obtained from published sources (Brady et al., 2008).

#### Training phase

Subjects learned to look for four different sets of unique real-world objects of varying set size (1, 2, 4, or 6) on four different colored background contexts. The learning phase had them search for these objects repeatedly across 120 trials with each context. For example, they might learn that a red context was associated with a football while a green context was associated with a hat and a jacket, and so on (Figures 1a & 1c). The assignment of these individual objects to different set size conditions was randomized across subjects. Subjects indicated that one of the 6 objects in a search array matched the target(s) by responding present by pressing a button on a keyboard with their right index and they responded absent by pressing a key with their middle finger. Subjects were explicitly instructed that the searched-for objects would remain the same for each background color, except for the non-repeating-context condition, so that they should do their best to learn to look for these objects in their color context.

Each trial started with a color-background context, which stayed on the screen until the end of the trial (a square with the width of 13.42° visual angle). Then, 1000ms after trial onset, a specific set of unique real-world objects appeared for 500ms, followed by a blank delay of 2000ms. For each subject, object locations for each set size were randomly drawn from 6 locations around the fixation point (the eccentricity of 2.25° visual angle; 0, 60, 120, 180, 240, or 300° angular position). The object locations for individual set size conditions were the same across trials for each subject. These objects were presented on top of small white squares (width = 2.82° visual angle). The unfilled locations were also presented with these small white squares as placeholders so that the spatial extent of overall visual input across different set sizes was equated.

After the delay period, subjects saw a search array of 6 objects placed in front of the white square placeholders, at an eccentricity of 4.38° from the fixation (width = 3.69° visual angle). Object locations were randomly drawn from 6 locations (30, 90, 150, 210, 270, or 330° angular position). On target-present trials (50% of the trials), one of the 6 objects matched one to the objects presented before the delay period (referred to as a target) and the other 5 objects were distractors randomly selected from a pool of 900 unique objects. In the target-absent trials, all 6 objects were distractors, randomly selected from the same object pool. The search array appeared for 1500ms and subjects were instructed to respond as quickly and accurately as possible. The offset of the search array was then followed by a blank interval of 300ms and then feedback at the fixation dot for 200ms (“$” for correct responses and “X” for incorrect responses or responses slower than 1500ms). Trials were separated by inter-trial-intervals (ITIs) randomly drawn from a uniform distribution of 500-700ms. In addition, we interleaved these context-learning trials with non-predictive-context trials, in which we presented a fifth colored-background context followed by the possible target objects that never repeated across 120 trials (Figure 1b). The object set size also varied (1, 2, 4, or 6 objects; 30 trials for each set size). These non-repeated objects were pseudo-randomly selected from a different pool of 390 unique objects.

Overall there were 5 different color background contexts (i.e., red, green, blue, purple, and brown at photometric isoluminance: 9.14 cd/m^2^). The mapping between these color contexts and set size conditions were randomized across subjects to reduce confounds driven purely by physical differences between different colors. The training phase consisted of 15 blocks (600 trials in total, lasting ∼1.5-2 hours). Each block contained 40 trials (8 trials for each context).

#### Test phase

Five minutes after the training phase, subjects entered the test phase where they performed the visuals search task, without target cues, simply using the background color to determine if one of the targets from the set was present. The task during this phase was similar to that in the training phase except that the searched-for object(s) were not physically present. Subjects saw a blank color background context for 2000ms, followed by a search array. The distractors in the search array during the test phase were randomly selected from an image pool of 900 objects, different from the pool used during the training phase. This was done to prevent the learning of distractors across the two phases. The test phase consisted of 15 blocks (600 trials in total, lasting ∼1-1.5 hours). Each block contained 40 trials (8 trials for each context). For both training and test phases, subjects were asked to fixate the dot at the center of the screen and to minimize blinks, eye movements, and head movements, while performing the task. Overall, the experimental protocol including task instruction, EEG preparation, training, and testing lasted approximately 4-4.5 hours.

### Behavioral analysis

First, we sorted hit, false alarm, and RTs from correct trials (i.e. correct RTs) of individual subjects into different data bins according to their set size (1, 2, 4, versus 6), trail repetition (repeated versus non-repeated non-predictive-context), and phases(training versus test). Then, we computed target sensitivity (d’) during the training and test phases using the following equation: d’ = Z(hit) – Z(false alarm), where Z(x) is the inverse function for the cumulative distribution function of the Gaussian function. We also calculated averaged correct RTs for individual data bins.

To test learning effects from trial repetition during the training phase, we used two-way repeated-measures with within-subject factors of trial repetition (repeated versus non-repeated trials) and set size (1, 2, 4, versus 6) on the d’ and RT data. Then, we tested learning effects from trial repetition across the training and test phases using two-way repeated-measures analysis of variance (ANOVA) with within-subject factors of phase (training versus test) and set size (1, 2, 4, versus 6) on the d’ and RT data obtained from repeated trials (i.e., learned context). Finally, to examine the influence of trial repetition during the training phase on the retrieval performance during the test phase, we performed two-way repeated-measures analysis of variance (ANOVA) with within-subject factors of trial repetition (repeated versus non-repeated trials) and set size (1, 2, 4, versus 6) on the d’ and RT data in the test phase.

### EEG data acquisition

We recorded EEG data from the 10-20 electrode sites Fz, Cz, Pz, F3, F4, C3, C4, P3, P4, PO3, PO4, O1, O2, T3, T4, T5, and T6, and a pair of custom sites OL and OR, which were halfway between O1 and T5 and halfway between O2 and T6, respectively. These EEG data were referenced online to the right mastoid. Horizontal eye movements were monitored via a pair of external electrodes affixed ∼1cm lateral to the outer canthi of the left and right eyes. Blinks and vertical eye movements were tracked using an electrode placed below the right eye. The impedance of each electrode was kept below 3 KΩ. We amplified the EEG data with a gain of 20,000 using an SA Instrumentation amplifier with a bandpass filter of 0.01– 100 Hz and digitized at 250 Hz.

### EEG preprocessing and ERP analysis

We preprocessed the EEG data using custom MATLAB scripts and EEGLab11.0.3.1b (Delorme and Makeig, 2004). First, we re-referenced the EEG data offline to the average of the left and right mastoid electrodes. Next, we filtered the data using 0.25-Hz high-pass and 55-Hz low-pass Butterworth filters (3rd order). Next, we segmented the continuous EEG data into epochs extending from 2000ms before to 7000ms after trial onset. We then performed independent component analysis to remove prominent eye blinks (Makeig et al., 1996) and used threshold rejection and visual inspection to reject trials containing residual eye movements, muscle activity, drifts, and other artifacts. This artifact rejection protocol resulted in the removal of 19.13% ± 7.18% SD of trials across 36 subjects.

In the main text, we focused the EEG analysis on the test phase, where subjects had to use the learned color background context to perform a visual search task, however, we also present the ERP results during the training phase in Supplementary Figure 3. First, we sorted the data into 5 experimental bins: set size conditions 1, 2, 4, and 6 plus the non-predictive-context condition where the fifth color context was associated with a different target set on each training trial. We then time-locked the data to the context onset and the search onset and computed the mean values of the EEG signals to obtained the event-related potentials (ERPs). Following previous studies, we focused our analyses on 3 ERP components: the posterior delay activity (pDA), N2pc, and P3 components, which we used to index the holding of representations in working memory, target selection, and target-related decision-making processes, respectively(Eimer, 1996; Hillyard et al., 1972; Woodman and Luck, 1999). We focused our analyses on the electrode site(s) that each of these components is maximal after verifying that the distribution of voltage was similar to these previous reports. The mean amplitude the pDA was computed from 400-900ms after the context onset over the Pz electrode. The N2pc was obtained from 200-300ms after the search onset and was computed by subtracting the ERPs ipsilateral to the target from those contralateral to the target over the T3, T4, T5, and T6 sites. The mean amplitude of the P3 was calculated from 300-900 after the search onset over the Cz. To plot the ERP data, we added an additional filter of 25Hz-lowpass filter. However, all reported statistical results were based on 0.25-55Hz filtered data (see similar methods in Hickey et al., 2010; Itthipuripat et al., 2015; Rungratsameetaweemana et al., 2018).

To test for set size effects on the mean amplitude of the pDA component, we used a one-way repeated-measures ANOVA with set size as a within-subject factor (1, 2, 4, versus 6). Post-hoc pair-wise t-tests were then used to examine differences in the pDA amplitude across set size. Multiple comparisons were corrected using the Holm-Bonferroni method. Lastly, we used 4 planned pair-wise t-tests to compare the mean amplitude of the pDA from each set size to the mean amplitude in the non-predictive-context condition. This was done to test if the pDA modulation reflected content-specific information pulled from LTM or merely indexed general task difficulty. We further tested the relationship between our WM activity and visual search performance by examining individual differences in the pDA. To do so, we first computed the differences between the pDA amplitudes for individual set size and the pDA amplitude in the non-predictive-context condition. Then, we computed the pair-wise linear correlation coefficients between the pDA amplitude differences and the mean behavioral d’ values obtained from the test phase.

For the N2pc and P3 components, we also performed one-way repeated-measures ANOVAs with set size as a within-subject factor. Post-hoc pair-wise t-tests were used to examine differences in the mean amplitudes of these components across all set size conditions, and multiple comparisons were corrected using the Holm-Bonferroni method. Finally, we use planned pair-wise t-tests to compare the mean amplitudes of the N2pc and P3 from each set size to those measured in the non-predictive-context condition.

### Alpha suppression analysis

We decomposed single-trial EEG data using spectrogram.m in MATLAB with a fixed window size of 400ms and a window overlap of 360ms to obtain Fourier coefficients at 8-13Hz in 1Hz steps. Next, we computed the amplitude of these Fourier coefficients, baseline-corrected from −600 to −200ms, and divided the resultant spectral difference by the baseline activity for each frequency and multiplied by 100. These steps yielded the percentage change in alpha amplitude, which was then averaged over 8-13Hz from the posterior occipital sites (PO3/PO4). To examine set size effects on alpha amplitude during the delay period, we computed the average of alpha amplitude across 600-1400ms after context onset and used a one-way repeated-measures ANOVA with set size as a within-subject factor. To examine set size effects on alpha amplitude during the search period, we computed the average of alpha amplitude across 600-1400ms after search onset and used a one-way repeated-measures ANOVA with set size as a within-subject factor. Because this ANOVA revealed a significant set size effect, we performed post-hoc pair-wise t-tests to examine differences in the mean amplitudes of these components across all set size conditions, and multiple comparisons were corrected using the Holm-Bonferroni method. Finally, we used 4 planned pair-wise t-tests to compare alpha amplitude from each set size to the amplitude measured in the non-predictive-context condition.

## Supplementary Information

**Figure S1.**
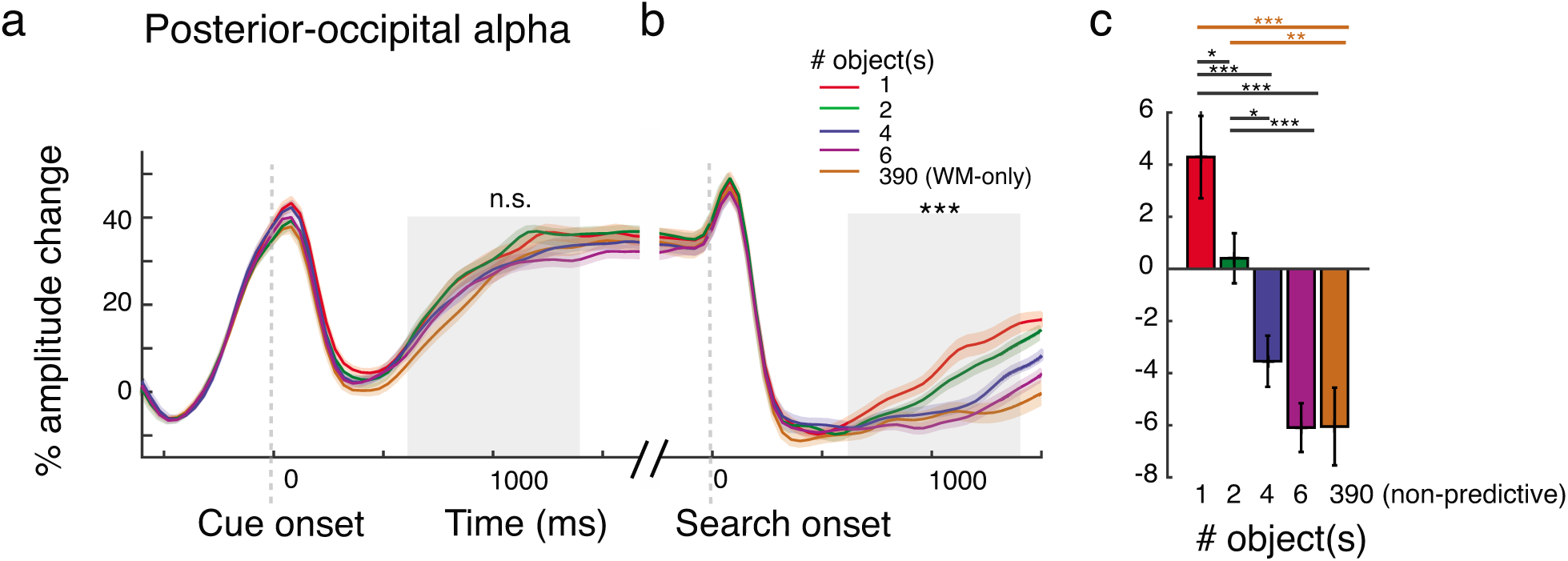
Alpha results from the training phase. (a) During the delay period, we did not observe any set size modulations of alpha suppression (testing across the 4 set sizes from 600-1400ms after context onset: F(3, 105) = 1.77, p =0.1583; including the non-predictive-context condition: F(4, 140) = 2.11, p =0.0826). (b-c) However, we observed an increase in the magnitude of alpha suppression as function of memory set size from 600-1400ms after the search onset (F(3, 105) = 13.22, p =1.17 × 10^−7^). Unlike the pDA result (Figure 3a), the magnitude of alpha suppression in the non-predictive-context condition was higher than those in set size 1 and 2 conditions (t(35)’s = 3.77 and 3.07, p’s = 0.0060 and 0.0041, respectively) and was comparable to the magnitude of suppression with set sizes 4 and 6 (t(35)’s = 1.21 and −0.03, p’s = 0.2332 and 0.9789). Together, these results are consistent with previous reports showing that alpha-band activity does not track the objects representations stored in WM, and may index the storage of purely spatial information, as well as general task demands(Bae and Luck, 2017; Bosman et al., 2012; Foster et al., 2017b, 2017a, 2016; Foxe and Snyder, 2011; Foxe et al., 1998; Fries, 2001; Fries et al., 2008; Fukuda and Woodman, 2017; Klimesch et al., 2007; van Moorselaar et al., 2017; Nelli et al., 2017; Rihs et al., 2007; Rungratsameetaweemana et al., 2018; Sauseng et al., 2005; von Stein et al., 2000; de Vries et al., 2017). (c) Graph of the alpha data shown in (b) averaged across a 600-1400ms window after search onset. *** in (b) show a significant set size effect in the alpha band averaged across the shaded areas via a repeated-measured one-way ANOVA with p’s <0.001. Black * and *** in (c) show significant differences from post-hoc pair-wise t-tests with p < 0.05 and p < 0.001, respectively (Holm-Bonferroni corrected). Brown ** and *** in (c) show significant differences from planned pair-wise t-tests between non-predictive-context trials and trials with learned contexts with p < 0.01 and p < 0.001, respectively. All shaded areas around alpha traces and all error bars represent ±1 within-subject SEM.

**Figure S2.**
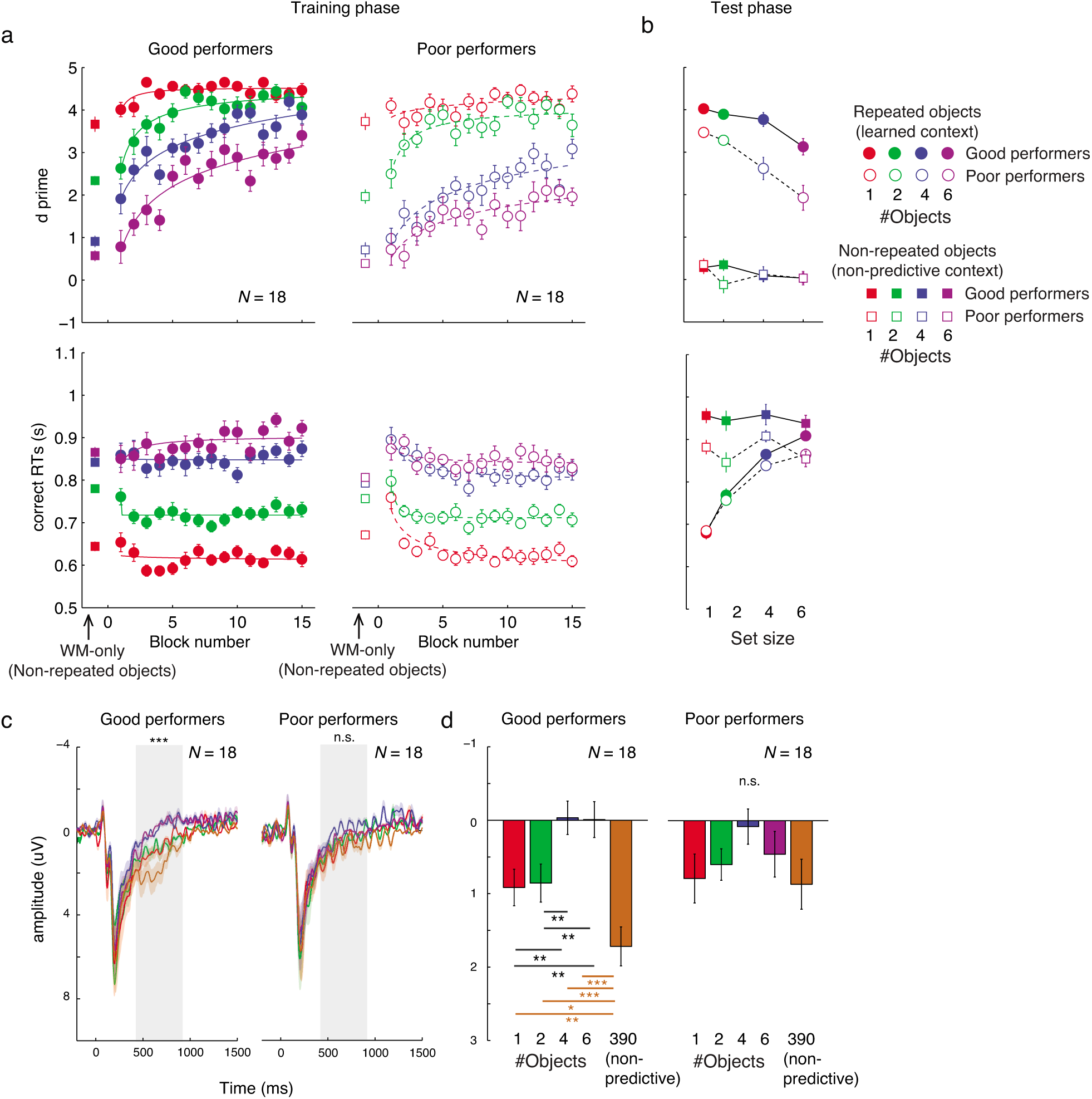
The behavioral and pDA data averaged separately using a median split between good and poor performers. The data were median split by the behavioral performance during the test phase. Specifically, we computed the difference in d’ values at large set sizes (averaged across set sizes 4 and 6) and d’ values at small set sizes (averaged across set size 1 and 2). This created two groups: good and bad performers with the former group having relatively smaller differential d’ values between high (4 and 6) vs low set sizes (1 and 2) than those in the latter group. (a) During the training phase, good performers learned much faster than poor performers especially for set sizes 4 and 6. (b) As expected given the sorting criterion, during the test phase the good performers performed better than poor performers when encountered learned contexts (i.e., repeated objects) especially for set sizes 4 and 6. However, both groups performed close to chance (d’ ∼ 0) in the non-predictive-context condition. (c-d). Good performers exhibited a robust and highly significant set size modulation in the pDA amplitude (400-900ms after context onset: F(3, 105) = 13.79, p ≤ 1.00 × 10^−15^). Similar to the main result in Figures 3.a.i-ii, the pDA amplitude increased as the number of to-be-searched-for objects increased (post-hoc parried t-tests showed significant differences between set sizes 1 and 4, between set sizes 1 and 6, between set sizes 2 and 4, and between set sizes 2 and 6: t(35)’s = 3.3152 to 2.9836, p’s = 0.0041 to 0.0010, Holm-Bonferroni corrected) and saturated at set size 4 (no difference between set sizes 4 and 6: t(35) = −0.08 with p = 0.9357). Importantly, when the number of to-be-searched-for objects was 390 (i.e., non-predictive-context trials), the pDA amplitude returned to the baseline activity level of set size 1 (planned paired t-tests showed significant differences between non-predictive-context trials and set sizes 2, 4, and 6: t(35)’s = −6.51 to −2.63, p’s ≤ 1.00 × 10^−15^ to 0.0177). While the pDA result in the poor performing group followed the data pattern observed in good performing group, the set size modulation in the poor performing group was weaker and not significant (testing across the 4 set sizes: F(3, 105) = 1.91, p =0.1402; including the non-predictive-context condition: F(4, 140) = 1.81, p =0.1366). *** in (c) shows a significant set size effects for the pDA data averaged across the shaded area via a repeated-measured one-way ANOVA with p <0.001. Black ** symbols in (d) show significant differences from post-hoc t-tests with p’s < 0.01 (Holm-Bonferroni corrected). Brown *, **, and *** in (d) show significant differences from planned t-tests between non-predictive-context trials and learned context conditions with p < 0.05, p < 0.01, and p < 0.001, respectively. All shaded areas around ERP traces and all error bars represent ±1 within-subject SEM. Note that we plotted negative up on all y-axes by electrophysiological convention.

**Figure S3.**
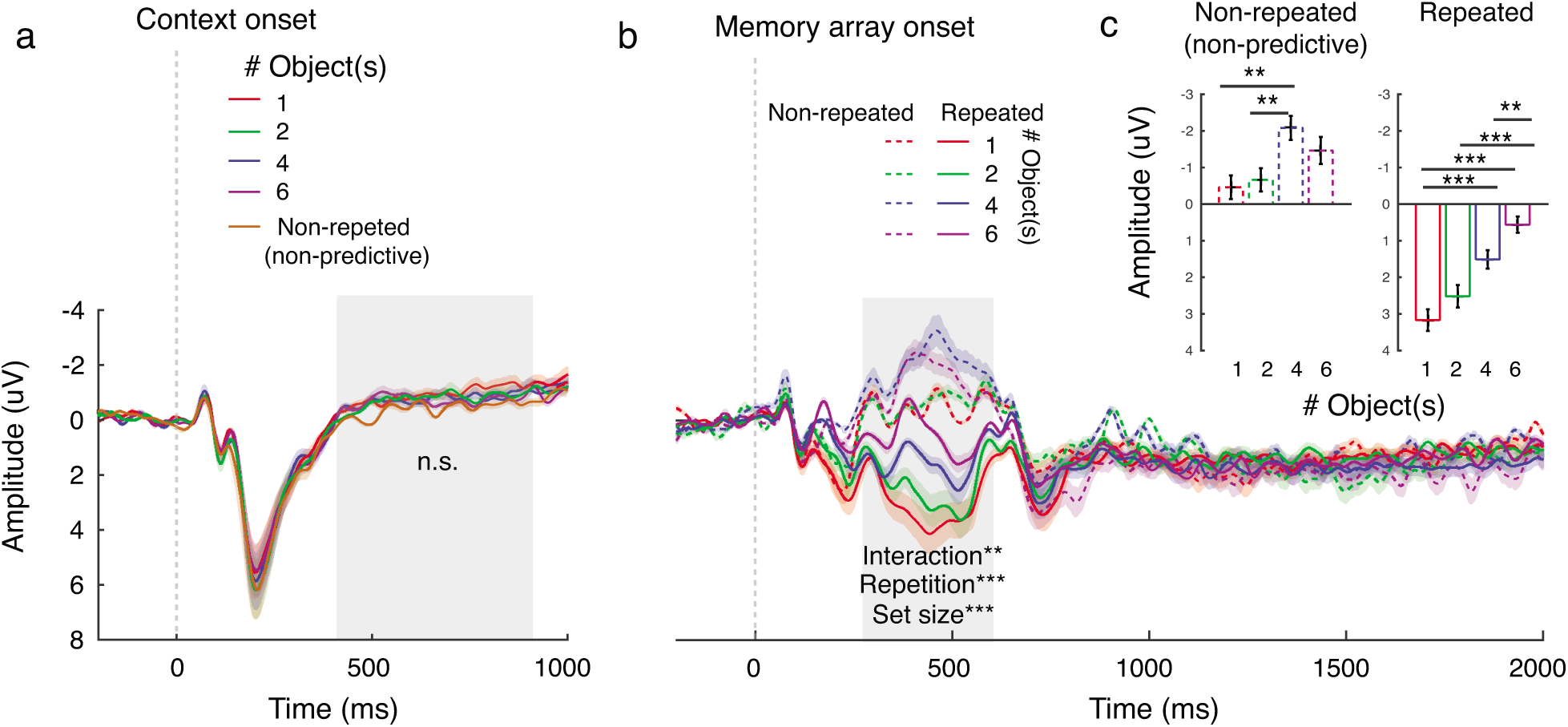
ERP results during the training phase. (a) Unlike the test phase, during the training phase there was no significant difference in the pDA amplitude across different color contexts from 400-900ms after the context onset (testing across the 4 set size conditions: F(3, 105) = 0.23, p =0.8762; including the non-predictive condition: F(4, 140) = 1.71, p =0.3254). Note that we averaged the non-predictive-context data across all set size because at this point in time, subjects did not know whether the subsequent memory array would contain 1, 2, 4, or 6 items. This null result confirms that the set size modulation of the pDA amplitude during the test phase selectively served LTM retrieval (Figure 3a), and it was not merely driven by physical stimulus differences between different color contexts or differences in learning difficulty across set sizes. (b-c) In contrast, we observed a robust set size effect on the posterior negative component over the Pz from 300-600ms after memory array onset (set size effect: F(3, 105) =13.26, p = 2.09 × 10^−7^). In the non-predictive-context condition, the amplitude of this negative potential increased as set size increased (set size 4 > set sizes 1 and 2: t(35)’s = 3.32 and 3.24 and p’s = 0.0021 and 0.0026, Holm-Bonferroni corrected) and reached asymptote at set size 4 (set size 4 = set size 6: t(35) = −1.33, p = 0.1929). Consistent with past reports(Reinhart and Woodman, 2014, 2015; Servant et al., 2018; Woodman, 2013), trial repetition induces robust amplitude changes in the opposite direction (more positive; repetition effect: F(1, 35) = 94.48, p =1.77 × 10^−11^) and these changes are larger for smaller set size conditions where learning is relatively easier (set size × repetition interaction: F(3, 105) = 4.71, p = 0.0040). This explains why an increase in negative potentials induced by increasing set size does not reaches asymptote at set size 4. Specifically, we found that the amplitude in set size 4 was higher than that in set size 1 (t(32) = 4.34, p =1.14 × 10^−4^) and the amplitude in set size 6 were higher than those in set sizes 1, 2, and 4 (t(32)’s = 6.75, 5.07, and 3.37; p’s = 8.09 × 10^−8^, 1.29 × 10^−5^, and 0.0018, Holm-Bonferroni corrected). (c) is the ERP data shown in (b) averaged across a 300-600ms window after memory array onset. ** and *** significant results with p < 0.01 and p < 0.001, respectively. All shaded areas around ERP traces and all error bars represent ±1 within-subject SEM. Note that we plotted negative up on all Y-axes.

**Table S1.** Post-hoc pair-wise t-test for Figure 3. *, **, and *** indicate significant differences with p < 0.05, p < 0.01, and p < 0.001, respectively (Holm-Bonferroni corrected).

| Comparisons | t-tests<br>(p values) |  |  |
| --- | --- | --- | --- |
|  | pDA | N2pc | P3 |
| 1 vs. 2 objects | 0.57<br>(0.5722) | -1.97<br>(0.0568) | 4.25**<br>(0.0002) |
| 1 vs. 4 objects | 3.71***<br>(0.0007) | -5.07***<br>(1.29x10<sup>-5</sup>) | 7.39***<br>(1.21x10<sup>-8</sup>) |
| 1 vs. 6 objects | 2.73**<br>(0.0097) | -4.56***<br>(0.0001) | 9.45***<br>(3.64x10<sup>-11</sup>) |
| 2 vs. 4 objects | 4.41***<br>(0.0001) | -1.63<br>(0.1118) | 4.27***<br>(0.0001) |
| 2 vs. 6 objects | 2.66*<br>(0.0118) | -2.79**<br>(0.0085) | 6.53***<br>(1.55x10<sup>-7</sup>) |
| 4 vs. 6 objects | -0.97<br>(0.3398) | -0.69<br>(0.4932) | 3.19**<br>(0.0030) |

## References

Atkinson, R.C., and Shiffrin, R.M. (1968). Human memory: A proposed system and its control processes. In The Psychology of Learning and Motivation, K. Spence, and J. Spence, eds. (Academic, New York), pp. 89–195.

Bae, G.-Y., and Luck, S.J. (2017). Dissociable Decoding of Spatial Attention and Working Memory from EEG Oscillations and Sustained Potentials. J. Neurosci. 38, 2860–17.

Bosman, C.A., Schoffelen, J.M., Brunet, N., Oostenveld, R., Bastos, A.M., Womelsdorf, T., Rubehn, B., Stieglitz, T., De Weerd, P., and Fries, P. (2012). Attentional Stimulus Selection through Selective Synchronization between Monkey Visual Areas. Neuron 75, 875–888.

Brady, T.F., Konkle, T., Alvarez, G.A., and Oliva, A. (2008). Visual long-term memory has a massive storage capacity for object details. Proc. Natl. Acad. Sci. 105, 14325–14329.

Brady, T.F., Störmer, V.S., and Alvarez, G.A. (2016). Working memory is not fixed-capacity: More active storage capacity for real-world objects than for simple stimuli. Proc. Natl. Acad. Sci. 113, 7459–7464.

Broadbent, D.H. (1975). The magic number seven after fifteen years. Studies in LongTerm Memory, eds Kennedy A, Wilkes A (Wiley, London), pp 3–18. In Studies in LongTerm Memory, A. Kennedy, and A. Wilkes, eds. (Wiley, London), pp. 3–18.

Bundesen, C., Habekost, T., and Kyllingsbæk, S. (2005). A neural theory of visual attention: Bridging cognition and neurophysiology. Psychol. Rev. 112, 291–328.

Chun, M.M., and Jiang, Y. (1998). Contextual Cueing: Implicit Learning and Memory of Visual Context Guides Spatial Attention. Cogn. Psychol. 36, 28–71.

Chun, M.M., and Phelps, E.A. (1999). Memory deficits for implicit contextual information in amnesic subjects with hippocampal damage. Nat. Neurosci. 2, 844–847.

Cosman, J.D., Arita, J.T., Ianni, J.D., and Woodman, G.F. (2016). Electrophysiological measurement of information flow during visual search. Psychophysiology 53, 535–543.

Cowan, N. (2001). The magical number 4 in short term memory. A reconsideration of storage capacity. Behav. Brain Sci. 24, 87–186.

Desimone, R., and Duncan, J. (1995). Neural mechanisms of selective visual attention. Annu. Rev. Neurosci. 18, 193–222.

Eimer, M. (1996). The N2pc component as an indicator of attentional selectivity. Electroencephalogr. Clin. Neurophysiol. 99, 225–234.

Foster, J.J., Bsales, E.M., Jaffe, R.J., and Awh, E. (2017b). Alpha-Band Activity Reveals Spontaneous Representations of Spatial Position in Visual Working Memory. Curr. Biol. 27, 3216–3223.e6.

Foster, J.J., Sutterer, D.W., Serences, J.T., Vogel, E.K., and Awh, E. (2017a). Alpha-Band Oscillations Enable Spatially and Temporally Resolved Tracking of Covert Spatial Attention.

Foster, X.J.J., Sutterer, D.W., Serences, J.T., Vogel, E.K., and Awh, E. (2016). The topography of alpha-band activity tracks the content of spatial working memory. J. Neurophysiol. 168–177.

Foxe, J.J., and Snyder, A.C. (2011). The role of alpha-band brain oscillations as a sensory suppression mechanism during selective attention. Front. Psychol. 2, 1–13.

Foxe, J.J., Simpson, G. V, and Ahlfors, S.P. (1998). Parieto-occipital ~10 Hz activity reflects anticipatory state of visual attention mechanisms. Neuroreport 9, 3929–3933.

Fries, P. (2001). Modulation of Oscillatory Neuronal Synchronization by Selective Visual Attention. Science (80-.). 291, 1560–1563.

Fries, P., Womelsdorf, T., Oostenveld, R., and Desimone, R. (2008). The Effects of Visual Stimulation and Selective Visual Attention on Rhythmic Neuronal Synchronization in Macaque Area V4. J. Neurosci. 28, 4823–4835.

Fukuda, K., and Woodman, G.F. (2017). Visual working memory buffers information retrieved from visual long-term memory. Proc. Natl. Acad. Sci. 114, 5306–5311.

Fukuda, K., Mance, I., and Vogel, E.K. (2015). Power Modulation and Event-Related Slow Wave Provide Dissociable Correlates of Visual Working Memory. J. Neurosci. 35, 14009–14016.

Fukuda, K., Kang, M.-S., and Woodman, G.F. (2016). Distinct neural mechanisms for spatially lateralized and spatially global visual working memory representations. J. Neurophysiol. 116, 1715–1727.

Giesbrecht, B., Sy, J.L., and Guerin, S.A. (2013). Both memory and attention systems contribute to visual search for targets cued by implicitly learned context. Vision Res. 85, 80–87.

Goldfarb, E. V., Chun, M.M., and Phelps, E.A. (2016). Memory-Guided Attention: Independent Contributions of the Hippocampus and Striatum. Neuron 89, 317–324.

Greene, A.J., Gross, W.L., Elsinger, C.L., and Rao, S.M. (2007). Hippocampal differentiation without recognition: An fMRI analysis of the contextual cueing task. Learn. Mem. 14, 548–553.

Hannula, D.E., and Ranganath, C. (2009). The Eyes Have It: Hippocampal Activity Predicts Expression of Memory in Eye Movements. Neuron 63, 592–599.

Heinen, K., Sagliano, L., Candini, M., Husain, M., Cappelletti, M., and Zokaei, N. (2016). Cathodal transcranial direct current stimulation over posterior parietal cortex enhances distinct aspects of visual working memory. Neuropsychologia 87, 35–42.

Hickey, C., Chelazzi, L., and Theeuwes, J. (2010). Reward Changes Salience in Human Vision via the Anterior Cingulate. J. Neurosci. 30, 11096–11103.

Hillyard, S., Squires, K., Baue, J., and Lindsay, P. (1972). Evoked potential correlates of response criterion in auditory signal detection. Science (80-.). 177, 1357–1360.

Itthipuripat, S., Ester, E.F., Deering, S., and Serences, J.T. (2014). Sensory gain outperforms efficient readout mechanisms in predicting attention-related improvements in behavior. J. Neurosci. 34, 13384–13398.

Itthipuripat, S., Cha, K., Rangsipat, N., and Serences, J.T. (2015). Value-based attentional capture influences context-dependent decision-making. J. Neurophysiol. 114, 560–569.

Itthipuripat, S., Cha, K., Byers, A., and Serences, J.T. (2017). Two different mechanisms support selective attention at different phases of training. PLoS Biol. 15, 1–38.

James, W. (1890). The Principles of Psychology (Henry Hold and Company, New York).

Klimesch, W., Sauseng, P., and Hanslmayr, S. (2007). EEG alpha oscillations: The inhibition-timing hypothesis. Brain Res. Rev. 53, 63–88.

Li, S., Cai, Y., Liu, J., Li, D., Feng, Z., Chen, C., and Xue, G. (2017). Dissociated roles of the parietal and frontal cortices in the scope and control of attention during visual working memory. Neuroimage 149, 210–219.

Luck, S., Vogel, J., and Edward, K. (1997). The Capacity of Visual Working Memory for Features and Conjuctions. Nature 390, 279–281.

van Moorselaar, D., Foster, J.J., Sutterer, D.W., Theeuwes, J., Olivers, C.N.L., and Awh, E. (2017). Spatially selective alpha oscillations reveal moment-by-moment trade-offs between working memory and attention. J. Cogn. Neurosci. 30, 256–266.

Nelli, S., Itthipuripat, S., Srinivasan, R., and Serences, J.T. (2017). Fluctuations in instantaneous frequency predict alpha amplitude during visual perception. Nat. Commun. 8.

Olivers, C.N.L., Peters, J., Houtkamp, R., and Roelfsema, P.R. (2011). Different states in visual working memory: When it guides attention and when it does not. Trends Cogn. Sci. 15, 327–334.

Preston, A.R., and Gabrieli, J.D.E. (2008). Dissociation between explicit memory and configural memory in the human medial temporal lobe. Cereb. Cortex 18, 2192–2207.

Rihs, T.A., Michel, C.M., and Thut, G. (2007). Mechanisms of selective inhibition in visual spatial attention are indexed by???-band EEG synchronization. Eur. J. Neurosci. 25, 603–610.

Rungratsameetaweemana, N., Itthipuripat, S., Salazar, A., and Serences, J.T. (2018). Expectations do not alter early sensory processing during perceptual decision making. J. Neurosci.

Sauseng, P., Klimesch, W., Stadler, W., Schabus, M., Doppelmayr, M., Hanslmayr, S., Gruber, W.R., and Birbaumer, N. (2005). A shift of visual spatial attention is selectively associated with human EEG alpha activity. Eur. J. Neurosci. 22, 2917–2926.

Squires, N.K., Squires, K.C., and Hillyard, S.A. (1975). Two varieties of long-latency positive waves evoked by unpredictable auditory stimuli in man. Electroencephalogr. Clin. Neurophysiol. 38, 387–401.

von Stein, A., Chiang, C., and Konig, P. (2000). Top-down processing mediated by interareal synchronization. Proc. Natl. Acad. Sci. 97, 14748–14753.

Summerfield, J.J., Lepsien, J., Gitelman, D.R., Mesulam, M.M., and Nobre, A.C. (2006). Orienting attention based on long-term memory experience. Neuron 49, 905–916.

Todd, J.J., and Marois, R. (2004). Capacity limit of visual short-term memory in human posterior parietal cortex. Nature 428, 751–754.

Tseng, P., Hsu, T.-Y., Chang, C.-F., Tzeng, O.J.L., Hung, D.L., Muggleton, N.G., Walsh, V., Liang, W.-K., Cheng, S. -k., and Juan, C.-H. (2012). Unleashing Potential: Transcranial Direct Current Stimulation over the Right Posterior Parietal Cortex Improves Change Detection in Low-Performing Individuals. J. Neurosci. 32, 10554–10561.

Vogel, E.K., and Machizawa, M.G. (2004). Neural activity predicts individual differences in visual working memory capacity. Nature 428, 748–751.

Vogel, E.K., McCollough, A.W., and Machizawa, M.G. (2005). Neural measures reveal individual differences in controlling access to working memory. Nature 438, 500–503.

de Vries, I.E.J., van Driel, J., and Olivers, C.N.L. (2017). Posterior α EEG Dynamics Dissociate Current from Future Goals in Working Memory-Guided Visual Search. J. Neurosci. 37, 1591–1603.

Wolfe, J.M. (2012). Saved by a Log: How Do Humans Perform Hybrid Visual and Memory Search? Psychol. Sci. 23, 698–703.

Wolfe, J.M., and Horowitz, T.S. (2004). What attributes guide the deployment of visual attention and how do they do it? Nat. Rev. Neurosci. 5, 495–501.

Woodman, G.F., and Luck, S.J. (1999). Electrophysiological m easurement of rapid shifts of attention during visual search. Nature 400, 867–869.

Woodman, G.F., Arita, J.T., and Luck, S.J. (2009). A cuing study of the N2pc component: An index of attentional deployment to objects rather than spatial locations. Brain Res. 1297, 101–111.

Woodman, G.F., Carlisle, N.B., and Reinhart, R.M.G. (2013). Where do we store the memory representations that guide attention? J. Vis. 13, 1–1.

Xu, Y., and Chun, M.M. (2006). Dissociable neural mechanisms supporting visual short-term memory for objects. Nature, 91–95. ns of electrophysiology. Vision Res. 80, 7–18.

## References for materials and methods

Brainard, D.H. (1997). The Psychophysics Toolbox. Spat. Vis. 10, 433–436.

Delorme, A., and Makeig, S. (2004). EEGLAB: An open source toolbox for analysis of single-trial EEG dynamics including independent component analysis. J. Neurosci. Methods 134, 9–21.

Hillyard, S., Squires, K., Baue, J., and Lindsay, P. (1972). Evoked potential correlates of response criterion in auditory signal detection. Science (80-. ). 177, 1357–1360.

Makeig, S. J. Bell., A., Jung, T.-P., and Sejnowski, T.J. (1996). Independent Component Analysis of Electroencephalographic Data. Adv. Neural Inf. Process. Syst. 8, 145–151.

Pelli, D. (1997). The VideoToolbox software for visual psychophysic:transforming numbers into movies. Spat. Vis. 10, 437–442.

## Supplementary information references

Foxe, J.J., Simpson, G. V, and Ahlfors, S.P. (1998). Parieto-occipital ∼10 Hz activity reflects anticipatory state of visual attention mechanisms. Neuroreport 9, 3929–3933.

Fries, P. (2001). Modulation of Oscillatory Neuronal Synchronization by Selective Visual Attention. Science (80-. ). 291, 1560–1563.

van Moorselaar, D., Foster, J.J., Sutterer, D.W., Theeuwes, J., Olivers, C.N.L., and Awh, E. (2017). Spatially selective alpha oscillations reveal moment-by-moment tradeoffs between working memory and attention. J. Cogn. Neurosci. 30, 256–266.

Reinhart, R.M.G., and Woodman, G.F. (2014). High stakes trigger the use of multiple memories to enhance the control of attention. Cereb. Cortex 24, 2022–2035.

Reinhart, R.M.G., and Woodman, G.F. (2015). Enhancing long-term memory with stimulation tunes visual attention in one trial. Proc. Natl. Acad. Sci. 112, 625–630.

Rihs, T.A., Michel, C.M., and Thut, G. (2007). Mechanisms of selective inhibition in visual spatial attention are indexed by ??-band EEG synchronization. Eur. J. Neurosci. 25, 603–610.

Servant, M., Cassey, P., Woodman, G., and Logan, G. (2018). Neural bases of automaticity. J. Exp. Psychol. Learn. Mem. Cogn. 44, 440–464.

Woodman, G.F. (2013). Viewing the dynamics and control of visual attention through the lens of electrophysiology. Vision Res. 80, 7–18.

